## Supplementary figures and images for "Loss of ErbB3 redirects Integrin β1 from early endosomal recycling to secretion in extracellular vesicles"

### Supplemental Figure 1

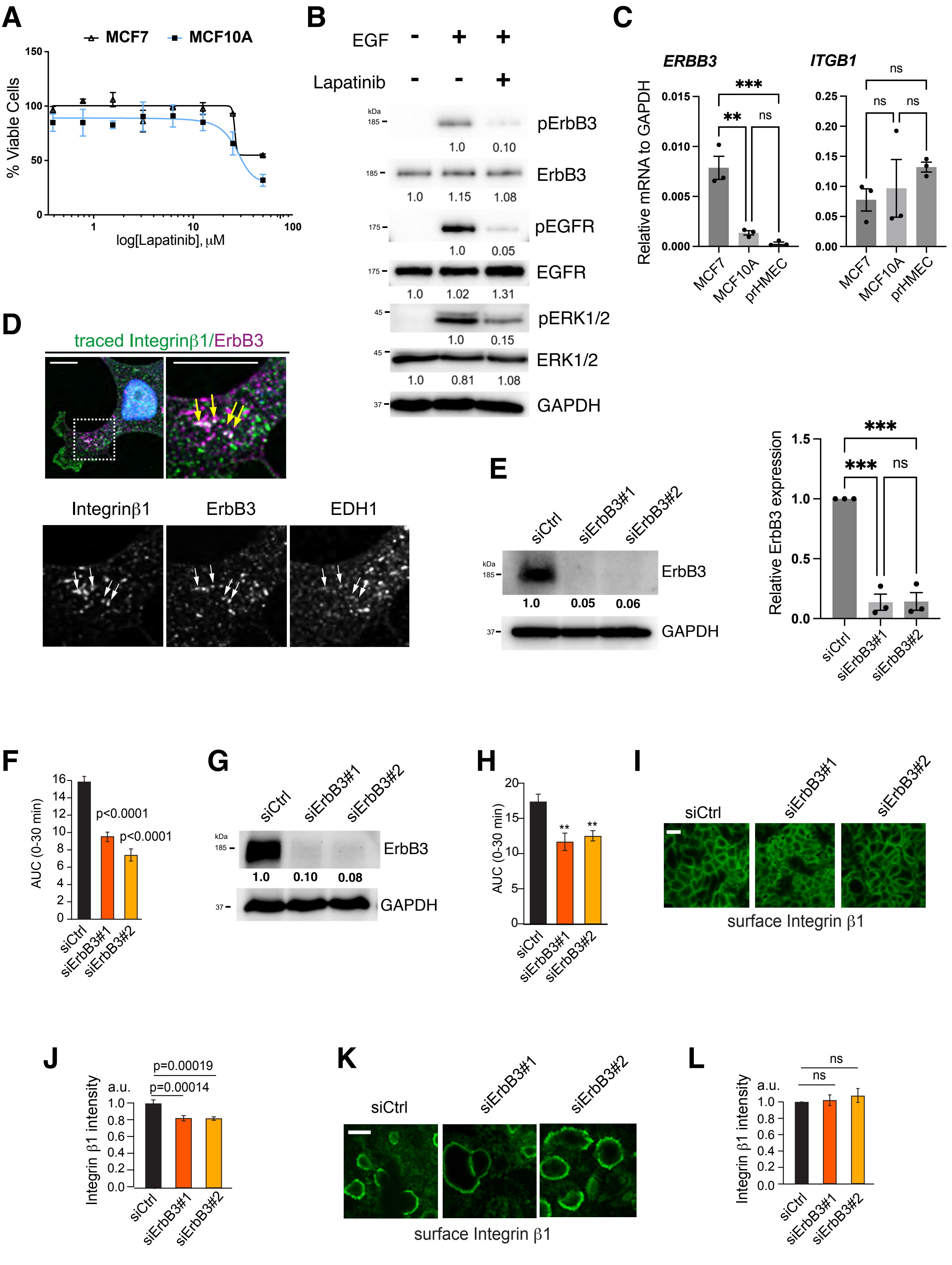


Suppl. Figure S1

### Supplemental Figure 2

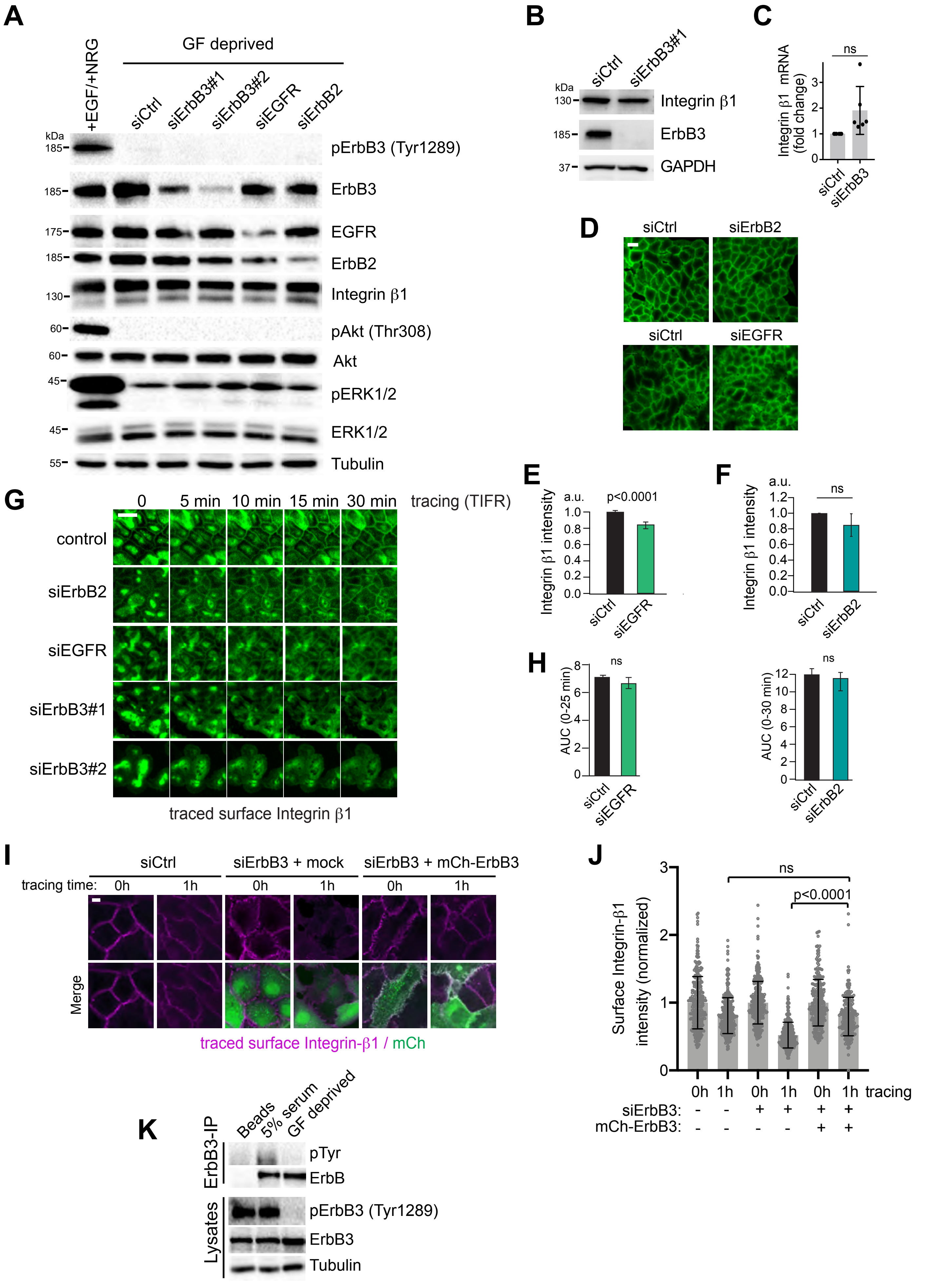


**Suppl. Figure S2**

### Supplemental Figure 4

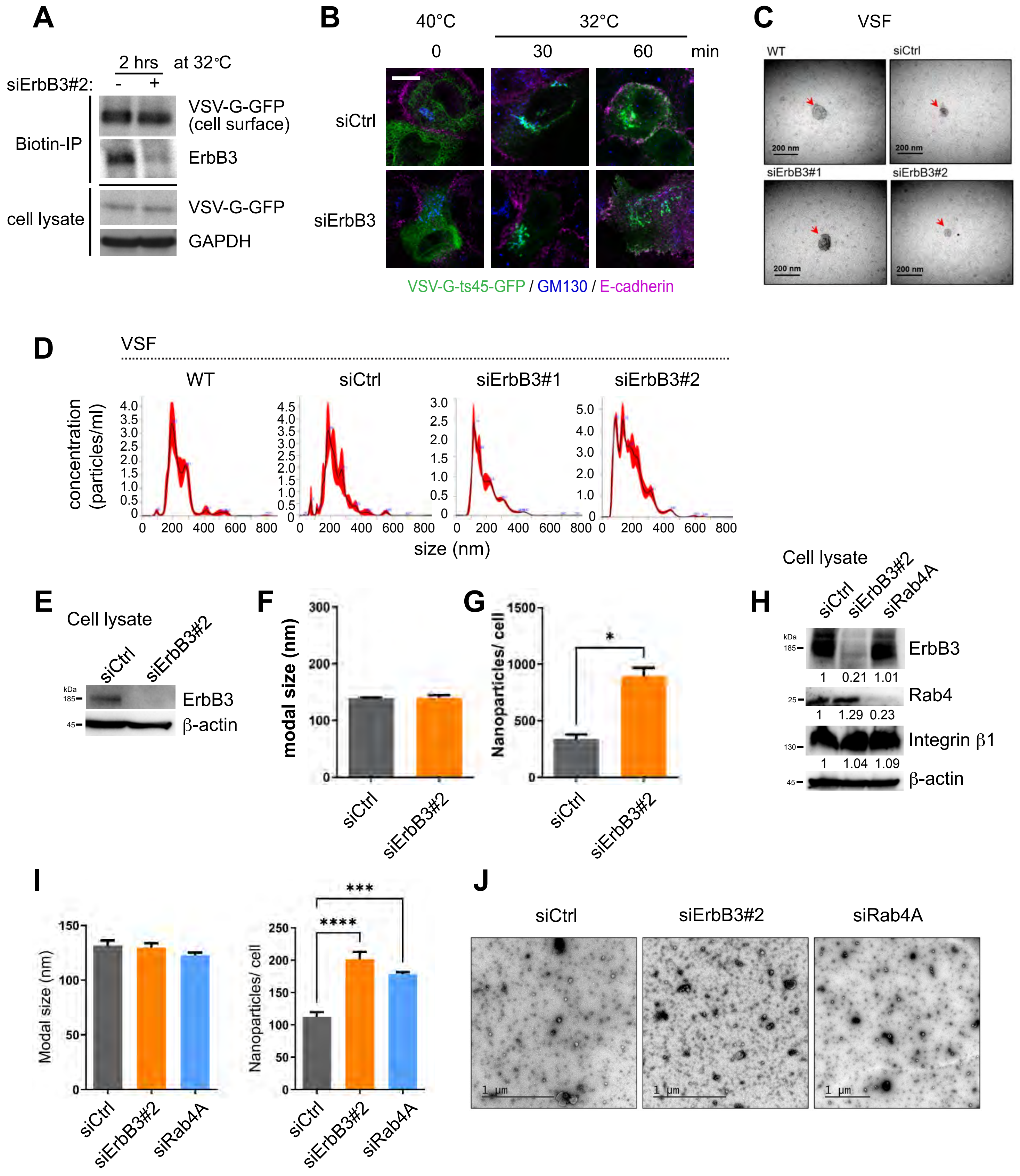


**Suppl. Figure S4**

### Supplemental Figure 5

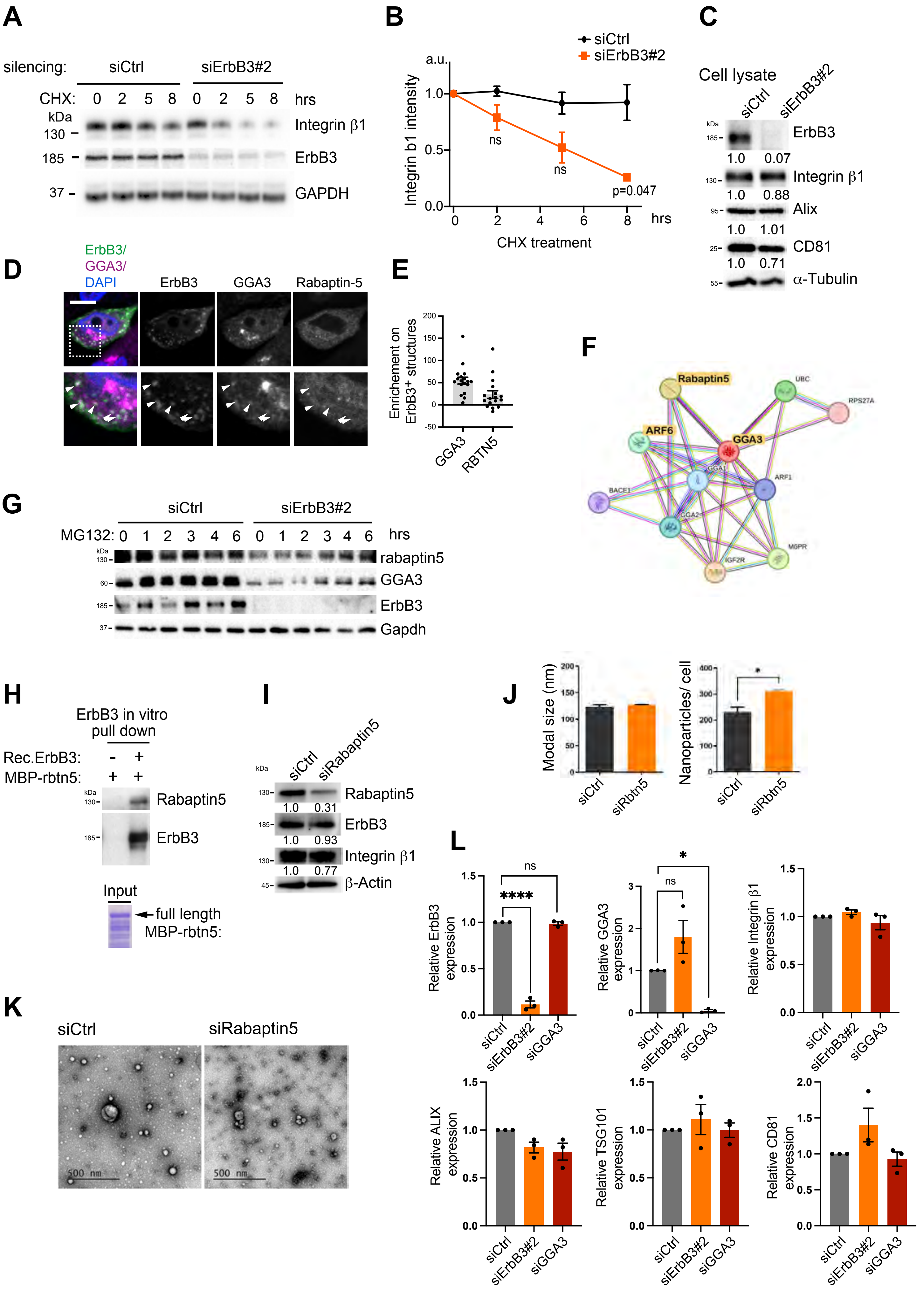


**Suppl. Figure S5**
