## Supplemental Figure 3 for "Loss of ErbB3 redirects Integrin β1 from early endosomal recycling to secretion in extracellular vesicles"

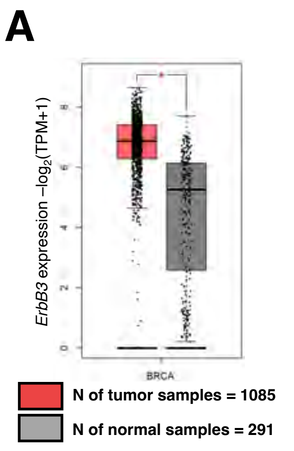


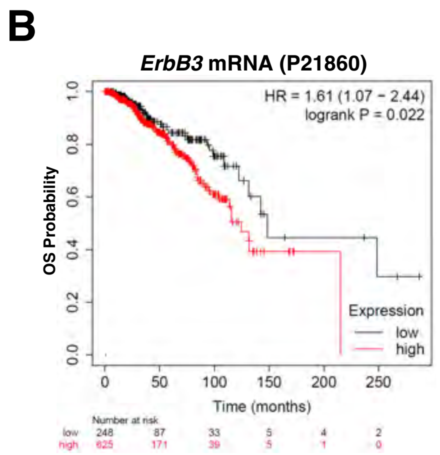


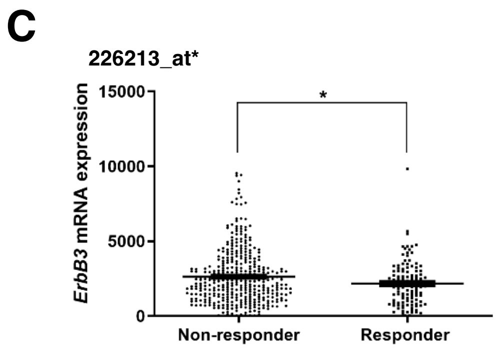


| Chemotherapy response | N of patients | Mean | SD | Median | Minimum | Maximum |
|---|---|---|---|---|---|---|
| Responder | 119 | 2174 | 1460 | 2097 | 26 | 9819 |
| Non-responder | 388 | 2631 | 1803 | 2279 | 27 | 9539 |

*p = 0.036

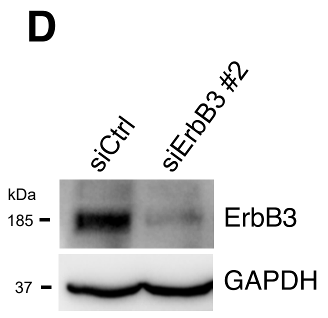


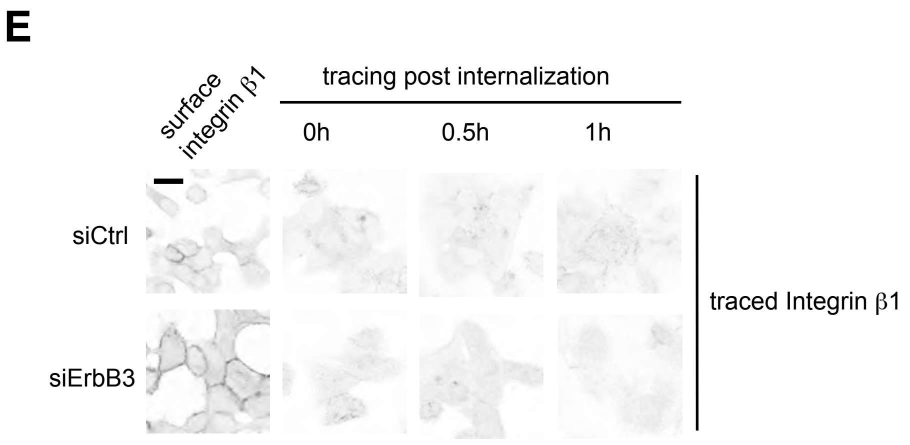


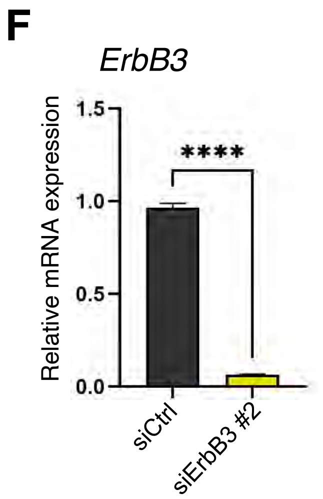


Suppl. Figure S3
